# Uncover the Mechanism of Nucleotide Import by HIV-1 Capsid

**DOI:** 10.1101/717173

**Authors:** Guang Song

## Abstract

In this work, we carry out a series of Brownian diffusion simulations to elucidate the nucleotide importing process through the hexamer pores of HIV-1 capsid. Our simulations reveal the mechanism by which deoxynucleoside triphosphates (dNTPs) diffuse through the arginine ring, the role of electrostatic potential within the pore in the importing process, and surprisingly, how IP6s and ATPs, though competing with dNTPs for binding at the arginine ring, end up accelerating the nucleotide import rate by thousands of folds so that it is sufficiently high to fuel the encapsidated DNA synthesis.

**IMPORTANCE:** Efficient nucleotide import is critical to fuel the reverse DNA synthesis that takes place within the capsid. However, the mechanism by which capsid imports the nucleotides is presently unclear. The current study through Brownian simulations shed insights into the process. A clear understanding of the mechanism of nucleotide import can be informative in developing drugs that target the process.

## 1 Introduction

Much progress has been made in understanding HIV-1 virus and its genome, structure, and life cycle since its first isolation and identification about three and half decades ago^1,2^.

In the area of drug development against the virus, since the first successful clinical trial on AZT (azidothymidine) in 1987^3^, over two dozens of FDA-approved HIV anti-viral drugs have been developed. These drugs function by interfering with the various stages of the viral life cycle, namely cell entry, membrane fusion, reverse transcription, integration to the host genome, maturation of new viral particles, and as such, they fall into several distinct families, namely, (1) entry inhibitors,(2) fusion inhibitors, (3) nucleoside-analog reverse transcriptase inhibitors (NRTIs) or non–nucleoside reverse transcriptase inhibitors (NNRTIs),(4) integrase inhibitors, and (5) protease inhibitors (PIs)^2,4^. Since presently no vaccine has been developed and there is no cure, these drugs, though presenting significant toxicity and side effects, have to be taken for a life time. Poor adherence leads to drug resistance and consequently there is a continuous need for new drugs, especially those that are long-acting and/or having fewer side effects.

The viral capsid, the protein shell that shields the viral genome upon its release into the host cell’s cytoplasm and whose atomic structure was determined recently^5^, plays significant roles in various steps of the viral life cycle^6^. It keeps the viral genome in a closed environment for efficient reverse DNA transcription and protects it from cytosolic nucleases such as cyclic GMP-AMP synthase (cGAS)^7^ and three-prime repair exonuclease 1 (TREX1)^8^. The capsid plays a key role also in nuclear entry through binding with nuclear targeting cofactors^9^. It has been long recognized as a potential drug target^9,10^. However, though a number of small molecule compounds have been shown to be good capsid inhibitors, there currently exists no FDA-approved HIV-1 anti-viral drugs in this family^11^. Fortunately, some of these capsid inhibitors, showing high potential to be effective against a broad range of HIV-1 strains and to be long-acting, are undergoing clinical trials^10,12^.

In a recent work, the capsid, which is composed over 200 hexamers and 12 pentamers, was proposed to import nucleotides for encaspidated DNA synthesis through its hexamer pores^13^. The authors identified a key structure feature, the positively-charge arginine ring (R18), in the bottle neck of the pore. The concentration of six like charges at the arginine ring creates a destabilizing effect on the capsid on one hand, but helps recruit the negatively-charged nucleotides on the other. One significance of the work is that if the pore indeed is the sole channel for nucleotide import, it could serve as another possible drug target on the capsid. Presumably some inhibitors could be designed to compete for nucleotide binding at the ring and thus cut off the supply line of nucleotides needed for DNA synthesis^13^.

This proposed import process however was challenged by a recent work by Mallery et al^14^ and many questions regarding the role of hexamer pores and the arginine ring were raised: If the role of the arginine ring is to facilitate dNTP import, how does it survive the competition of other polyanions such as ATP or IP6 that is either more abundant (ATP) or has a stronger binding affinity to the ring (IP6)? Can the pore distinguish between between ATP and dNTP? Are ATPs, whose concentrations are over 100 folds higher than dNTPs, competitors for dNTP recruitment through the pores? It is naturally expected that the presence of ATP should slow down the dNTP transport, and yet the authors found that “Surprisingly, even at a 100-fold excess of ATP over dNTPs, not only did ATP fail to inhibit ERT but it actually increased the quantity of measured transcripts”^14^. Why? ATP’s effect on ERT (encapsidated reverse transcription) was found to be dose-dependent and that “ATP only promoted ERT at concentrations of dNTPs *<* 100 mM”^14^. How to explain this dose-dependence? Moreover, if the pore is designed to facilitate the transport of compounds with negative charges, then it should be able import compounds with negative charges more efficiently than those without. Particularly, the negatively-changed nucleotide-based reverse transcription inhibitors (NRTIs) should be able to enter the capsid more easily than non-nucleoside reverse-transcriptase inhibitors (NNRTIs). However, the authors found that the pores were able to import both inhibitors equally well. Lastly, Mallery et al.^14^ found that polyanions such as ATPs and IP6s, when bound at the R18 ring, have a significant effect in stabilizing the capsid. Therefore, the arginine ring may have a dual role: one is to regulate the stability of the capsid through interacting with some pocket factor such as IP6^14^, the other is for nucleotide import. The two roles seem to be at odds with each other and needs to be reconciled, or, is there an alternative route into the capsid? The authors stated, “The fact that high concentrations of either ATP or IP6, which would be competitors with dNTPs for pore binding, increase rather than decrease ERT efficiency argues against the R18 pore being essential for nucleotide import.”^14^

In this work, we present a series of Brownian simulation studies that answer most of these questions. Particularly, we are able to reconcile the dual role of the arginine ring. We are able to explain why IP6/ATP, though being strong competitors with dNTP binding, still ends up accelerating dNTP import rate by thousands of folds and consequently the rate of ERT.

Our diffusion simulations are inspired by Kornberg and Levitt and coworkers’ diffusion model of the nucleotides^15^, in which the authors were able to use a diffusion model to understand how nucleoside triphosphates (NTPs) reach the active center of RNA ploymerase II through a funnel-shaped chamber and a narrow negatively-charge pore. The simulations provided significant insights into the process itself, the roles of several factors such as pore topography, electrostatics, and NTP interactions, and how the binding at the E site helps enhance the rate of transcription^15^.

In this work, we employ a similar Brownian diffusion model to understand the dNTP importing process through the hexamer pores of HIV-1 capsid. The topography of the hexamer pore also is funnel shaped, with a narrow opening in the center of the arginine ring. Our simulations reveal how nucleotides may diffuse through the arginine ring and surprisingly, how IP6 and ATP, though competing with nucleotides for binding at the arginine ring, accelerate the nucleotide import process. We provide also quantitative estimations of the diffusion rates and link them to the rates known from experiments.

## 2 Methods

### 2.1 The hexmer pore and it surrounding electrostatic potential

To understand the diffusion process of dNTPs through a capsid hexamer, we use one of the hexamer structures found in the atomic capsid structure 3J3Q^5^. Since our focus here is to understand how dNTP diffuse through the arginine ring and since most hairpins take an open conformation in general^16^, we select the seventh hexamer in 3J3Q since its hairpins are in a wide open conformation.

The electrostatic potential created by the hexamer is computed using the APBS software^17^, which solves the Possion-Boltzmann equation using the finite-element method. To this end, the convenient PDB/PQR web server^18^ is first used to generate a PQR (protein charge radius) file (hex7.pqr) and an input file (hex7.in), both of which are needed by the APBS software. The APBS software^17^ then computes the electrostatic potential around the hexamer and saves it in a discrete format, in a 3-D grid. The grid space is set by the PDB/PQR server^18^ and in our case is about 0.5 Å in all dimensions. The computation is done using an ionic strength (NaCl) of 0.15 M, a solvent dielectric constant of 78.54, and a protein dielectric constant of 2. We name this potential *ϕ*_*hex*_.

#### The electrostatic potential upon IP6 binding

In addition to the potential of the hexamer, we compute also the electrostatic potential of the hexamer in complex with IP6. IP6 is modeled as an ion with a net charge of −6 and a radius of 5 Å and is placed 5 Å above the arginine ring. This charge-radius information is attached to the hex7.pqr file and the potential is recomputed. We name this potential *ϕ*_*complex*_. The nucleotide on the other hand has a net charge of −4 units and is given a radius of 3.5 Å^15^.

### 2.2 The interaction between IP6 and dNTP

Most of our simulations are about the diffusion of a single dNTP particle under the influence of an electrostatic potential field within the hexamer pore, either with (*ϕ*_*complex*_) or without (*ϕ*_*hex*_) the influence of the IP6.

One of the simulations, on the other hand, is about the diffusion of both a dNTP and an IP6 (two moving particles). The interaction force **F**(**r**) (in the units of *Kcal/mol/*Å) between them takes the following form:

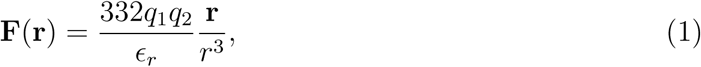

where *ϵ*_*r*_ is the dielectric constant and is set to 78.54 for solvent, **r** (r) is the distance vector (distance) between dNTP and IP6, *q*_1_ and *q*_2_ are the amount of charges of dNTP and IP6 respectively. The distance *r* is in Å. Note that this interaction implicitly takes into account the effect of solvent, but not the ions in the solvent. Ions, metal ions particularly, are likely to form complexes with dNTP and IP6 and thus may reduce the strength of their interaction. However, this effect is not considered here. The contributions of ions are considered though in the computations of potentials *ϕ_hex_* and *ϕ*_*complex*_.

### 2.3 Brownian Simulations

Inspired by the work by Kornberg, Levitt and co-workers^15^, we simulate the Brownian motions of dNTP (and IP6 too in one scenario) within the capsid hexamer pore in order to understand the importing process.

To facilitate the simulation and capture the key essence of the import process, we ignore the dynamics of the hexamer and treat it as a static structure. We model the influence of the hexamer on the diffusion of the nucleotide through its electrostatic potential. In other words, dNTP makes Brownian motions within the pore under the influence of the electrostatic potential of the hexamer.

The Brownian simulations are carried out by employing the following Langevin stochastic differential equations:

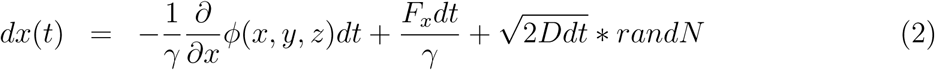

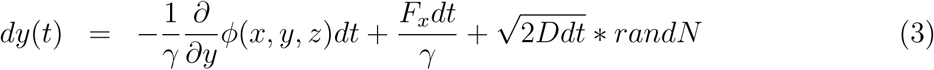

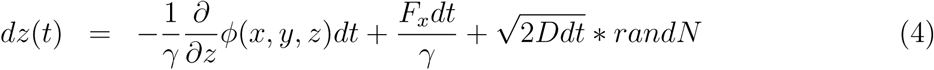

where (x(t), y(t), z(t)) represents the coordinates the simulated particle at time *t*. *ϕ* (either *ϕ*_*hex*_ or *ϕ*_*complex*_) is the electrostatic potential, D is the diffusion coefficient of the nucleotide and is set to be 75 *µm*^2^*/s*^15^, *γ* = *kT/D* where *k* is the Boltzmann constant and T is the temperature. *dt* is the time step is set to be 12.5 ps in all our simulations. randN represents a random number sampled from a standard normal distribution.

### 2.4 Boltzmann Distribution

Boltzmann distribution is a probability distribution according to the energy of states that a particle may occupy:

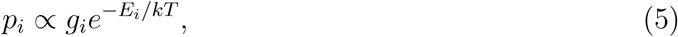

where *p*_*i*_, *g*_*i*_, *E*_*i*_ are the probability, degeneracy, and energy of state *i* respectively. *k* is the Boltzmann constant and T is temperature.

## 3 Results

Here, we perform Brownian simulations to understand the diffusion process of deoxynucleoside triphosphates (dNTPs) through the capsid hexamer’s R18 pore. Through performing realistic diffusion simulations, we aim to understand what determines the rate of dNTPs entering into the capsid. What is the role of IP6 or other nucleotides in dNTPs’ diffusion process? We aim to obtain a quantitative understanding of their roles and to link the simulation results with rates known from experiments.

### 3.1 The potential field in and around the pore

Fig. 1 shows the electrostatic potential within and around the hexamer pore in the side view (**A**) and the top view (**B**). The potential is computed using APBS software^17^ (see Methods section). The side view, where the ordinate axis is the pore axis, shows the potential within a vertical cross section of the hexamer that passes through the central pore axis. The arginine ring is near the origin, around which it is seen that there is a narrow passage that a nucleotide can pass through. The contour of the protein chains (only chains C and F are shown) are drawn using a probe sphere of radius 3.5 Å, the size of a nucleotide^15^. The size of this passage will vary as the protein chains fluctuate. There is a chamber right above that arginine ring as pointed out by Jacques et al.^13^, which is about 25 Å deep and has a volume of over 3,000 Å^3^ ^13^. There is an opening at the top the chamber at the location about 25 Å above the arginine ring. This is where the hairpins locate. As a result of the motions of the hairpins, the hairpin pore can open and close^13,16^. Presently in this figure the hairpins are in an open conformation. The space below the arginine ring represents the interior of the capsid. The potential contour (Fig. 1(A)) shows also that there is no other significant local energy minimum within the hexamer pore besides the one at the arginine ring. This means, once passing through the arginine ring, a nucleotide would diffuse freely into the interior of the capsid. The potential is computed using an isolated hexamer. The potential and the free space outside and around the hexamer should be ignored as it is inaccessible for hexamers within a capsid.

**Figure 1.**
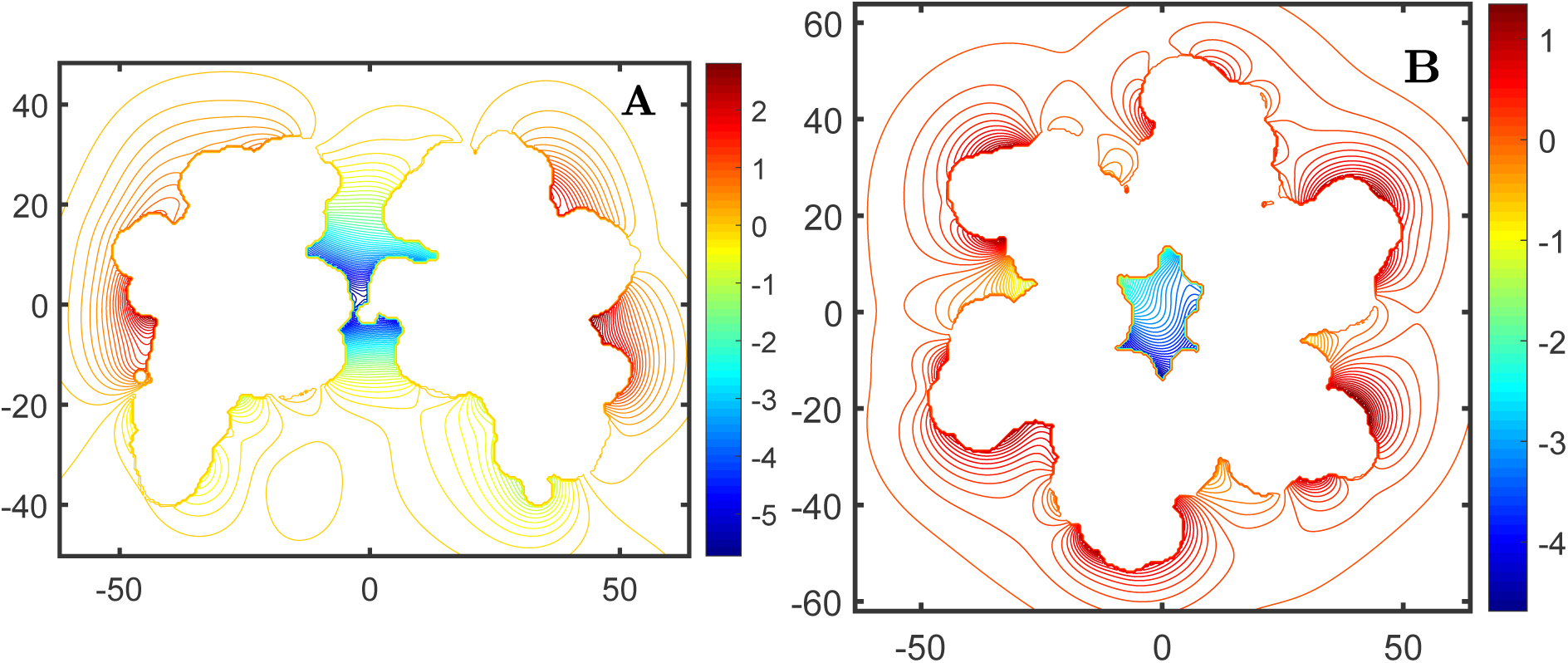
The electrostatic potential within and around the hexamer pore, in the side view (**A**) and the top view (**B**). The structure used here is the seventh hexamer in structure 3J3Q^5^.

Fig. 1(B) shows the potential within a horizontal cross section of the hexamer, in a plane that is 10 Å above the arginine ring, where the pore has the largest cross section. The central region of this top view represents a horizontal cross section of the chamber and shows the potential with it. Overall, the hexamer is positively charged within the pore (in blue), but negatively charged outside and around (in red).

The potential at the arginine ring is about −5.5 kT per one negative unit charge. For nucleotides that carry 4 negative charges, it would have to cross an energy barrier that is as high as 22 kT before entering capsid interior.

As will be shown next, the potential shown here, especially that within the pore, greatly influences the diffusion process of the nucleotides, in a fashion mathematically described in Eq. 4. Particularly, one significant consequence of this potential field is that it effects a deep energy well around the arginine ring (the blue region in the center of the figure) and thus a high concentration of nucleotides, which means that a given nucleotide, once bound to the arginine ring, will find it nearly impossible to diffuse away by itself, but rather be confined to a small region near the arginine ring. As will shown later, this extremely high concentration serves to greatly facilitate the diffusion of the nucleotide through the narrow hole/passage in the middle of the arginine ring.

### 3.2 Brownian motions and Boltzmann distributions

In the following, we perform Brownian motion simulations of the diffusion process of dNTP through the R18 pore. For easy visualization of the diffusion process, we first perform the diffusion simulation in 2-D, letting dNTP diffuse under the influence of the potential shown in Figure 1(A). We confine the dNTP diffusion to this 2D plane. We will then extend this simulation to the actual 3-D space.

Figure 2 shows the diffusion process of the dNTP. Under the influence of the potential, the nucleotide, initially placed at about 20 angstroms above the arginine ring, is quickly pulled towards the arginine ring and stays there through the rest of the simulation (black trajectory). The simulation lasts 10,000 steps. In contrast, also shown in the figure is the diffusion of dNTP without the influence of the electrostatic potential. As expected, it eventually diffuses away from the hexamer (trajectory in red).

**Figure 2.**
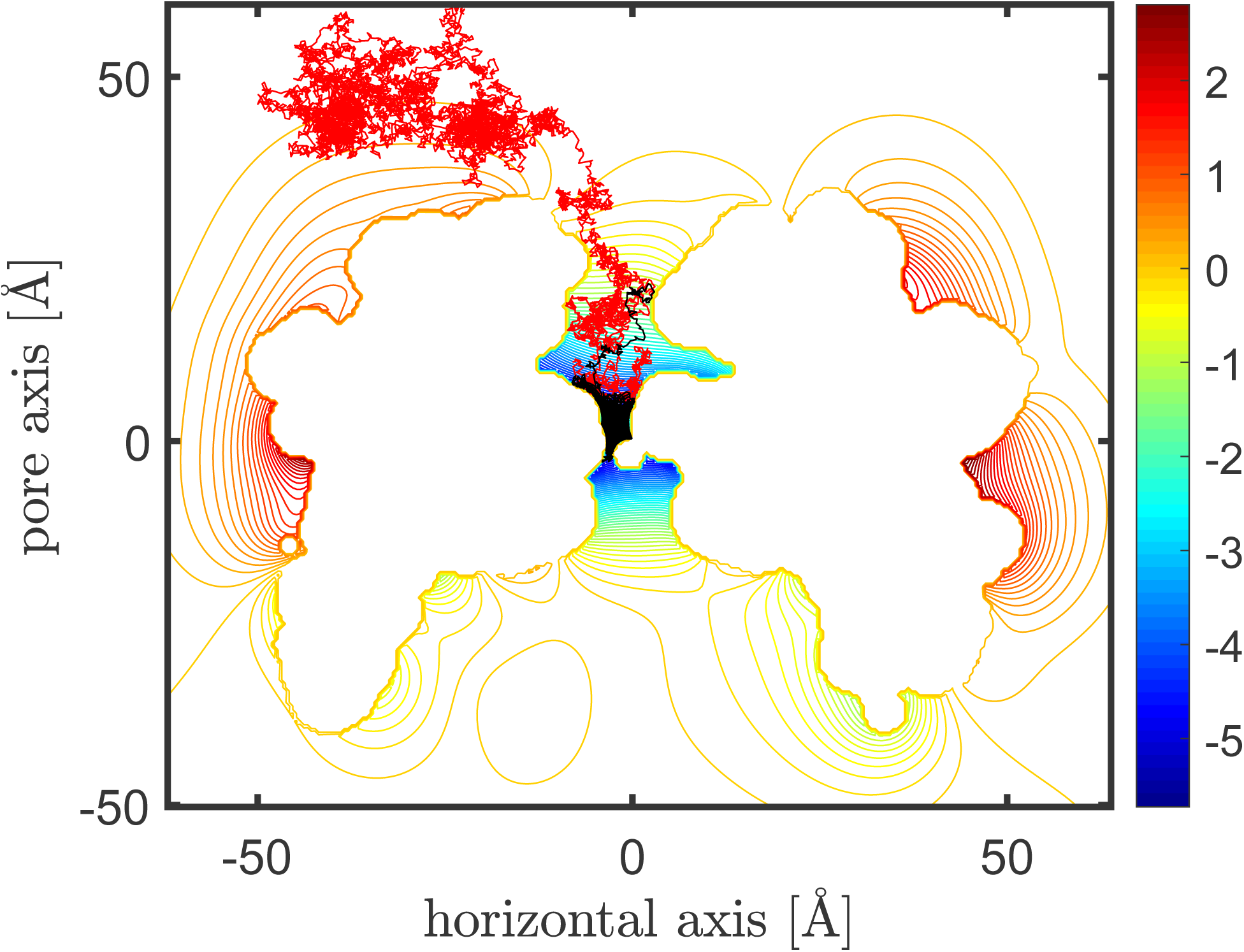
The diffusion trajectories of dNTP with (in black) and without (in red) the influence of the electrostatic potential within the pore. dNTP is quickly pulled towards the arginine and is trapped thereafter in the presence of the potential, but diffuse away in its absence.

Figure 3 shows the Brownian simulation in 3D. The nucleotide is initially placed at 20 Å above the arginine ring. As the same as in 2D (Fig. 2, the nucleotide is quickly pulled towards the arginine ring and get trapped in that region afterwards.

**Figure 3.**
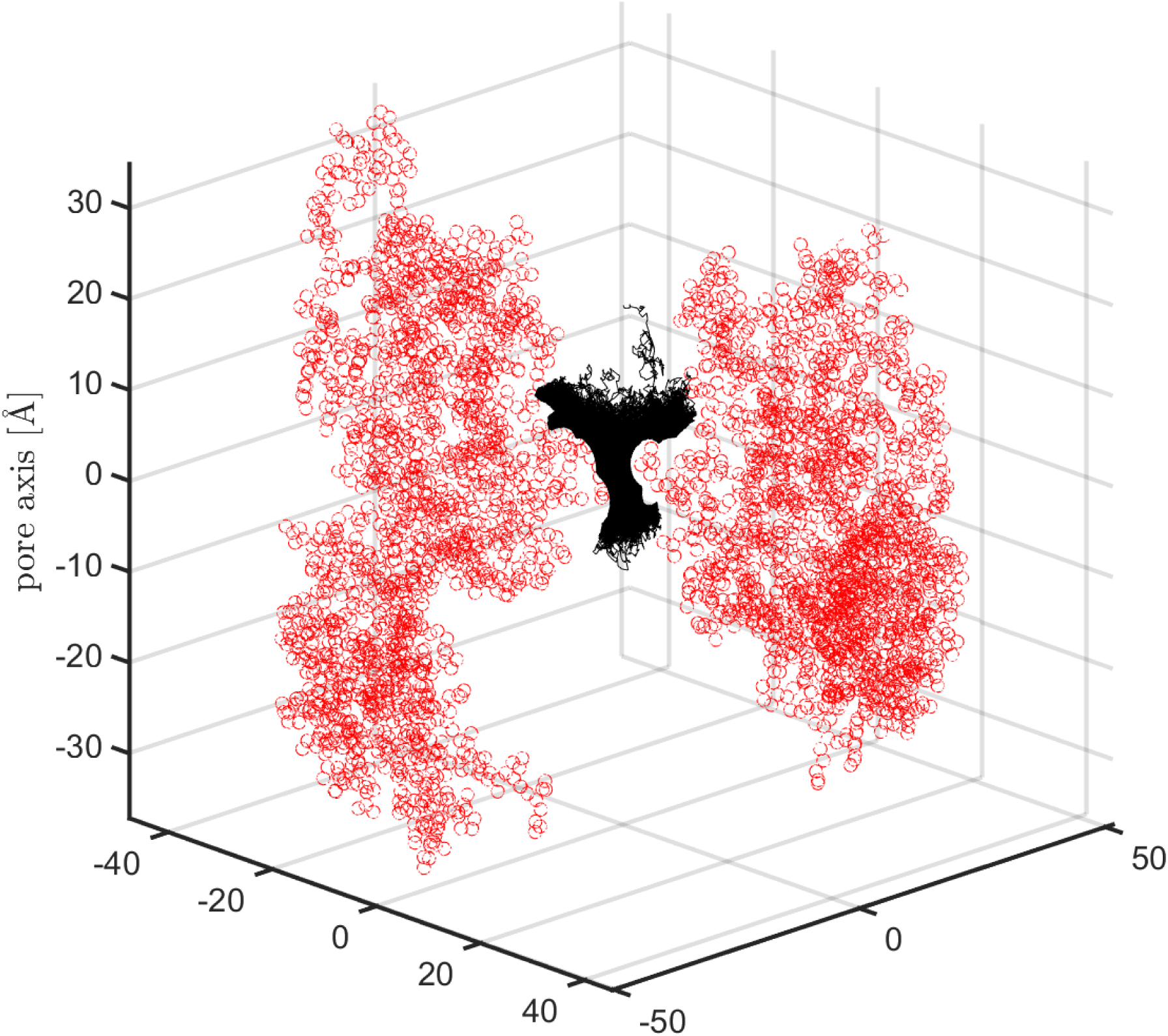
A 3D Brownian diffusion of dNTP within the pore. Similar to the 2D simulation shown in Fig 2, dNTP is quickly pulled towards the arginine ring and is trapped there.

To verify that the simulation results are correct, we plot the probability distributions of dNTP along the pore axis as a result of the simulations and compare them with the expected equilibrium Boltzmann distribution (Eq. 5, where *g*_*i*_ is set as 1). It is seen that there is a good agreement between the distributions from Brownian motions and the Boltzmann distribution. The small difference between the them reflects the actual topography of the chamber and variations of the potential within the chamber. The Boltzmann distribution (in red) does not take the geometry of the pore into account and uses only the potential along the pore axis. In all three distributions, it is clearly seen that there is a high concentration of dNTP at around the arginine ring, which is located near the origin.

### 3.3 High concentration around the arginine ring facilitates nucleotide translocation

The translocation of a nucleotide through the arginine ring could have been a challenging process. However, due to the deep potential well, a nucleotide, once bound to the arginine ring, is confined to a small vicinity around the arginine ring (see Figures 2, 3, and 4). Due to this high concentration at around the arginine ring, the diffusion or translocation through the arginine ring is greatly favored than retreating back or moving in any other direction. In other words, the potential in the chamber not only help recruit nucleotides by guiding them to where the opening of the ring is (which is at the center of the arginine ring), but also facilitates its diffusion through the narrow opening by confining the nucleotide within the vicinity of the opening.

**Figure 4.**
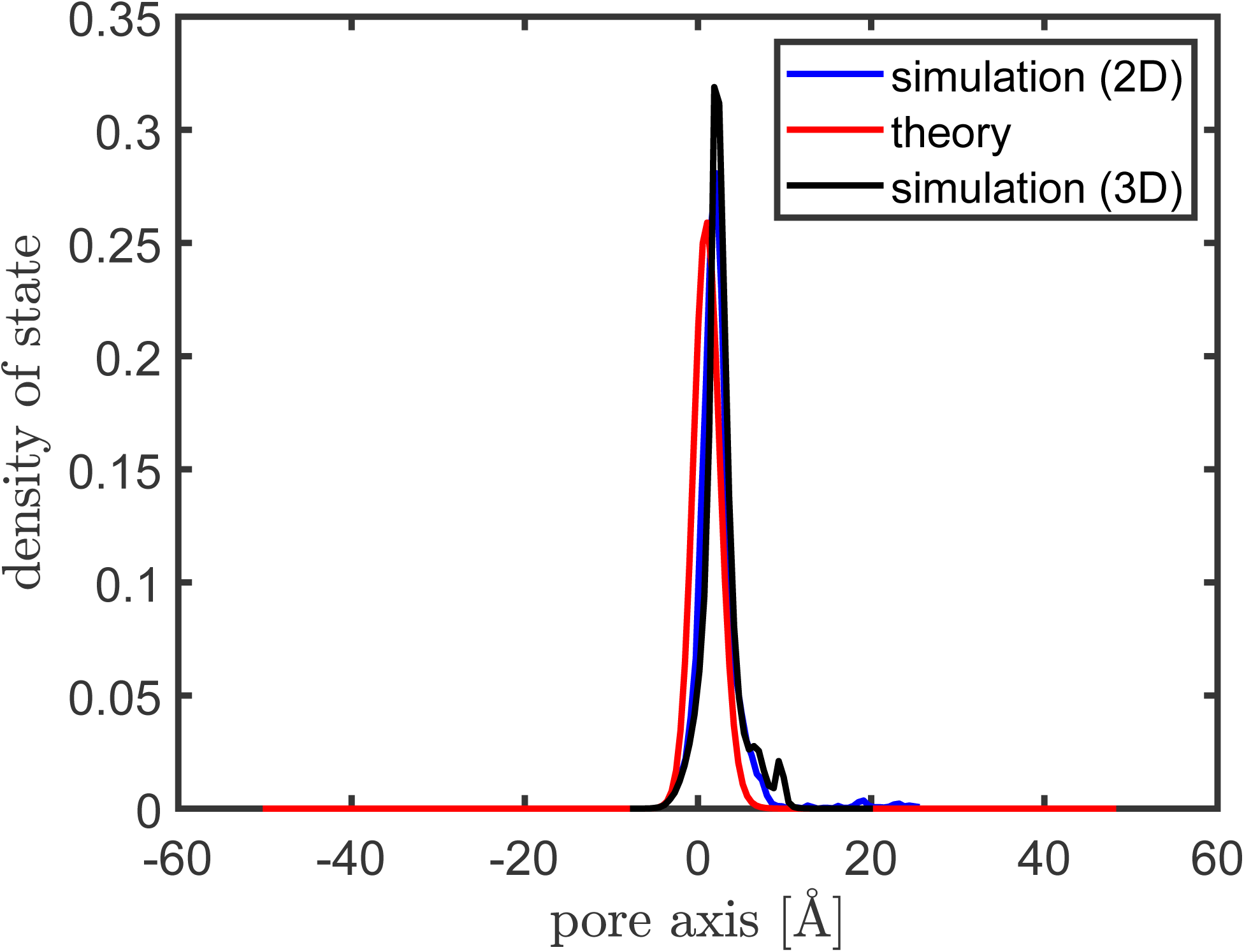
The probability distributions of dNTP along the pore axis according to a 2-D Brownian simulation (in blue), a 3-D Brownian simulation (in red), and Boltzmann distribution (in red). Both simulations, as expected, match extremely well with the equilibrium Boltzmann distribution. The difference reflects the effect of the pore topography.

Furthermore, the translocation should be facilitated by the swinging motions of the arginine side chains^16^. The arginine side chains are capable to bind and translocate the bound nucleotide for a downward movement for as much as 12 Å^16^.

### 3.4 The problem of release

While the deep potential well at the arginine ring plays an important role in attracting nucleotides and facilitating their crossing through the arginine ring, it poses also a problem: the problem of release. How can a bound nucleotide escape from such a deep well? Without outside help, the release rate would be too slow to fuel the reverse DNA synthesis going on inside the capsid.

Due to the attractive potential leading to the arginine ring, it was observed that nucleotides can arrive at the arginine ring at a diffusion-limited rate^13^, which is about 10^10^ *M^−^*^1^*s^−^*^1^ ^15^.

Kinetics of interaction by stopped-flow and steady-state affinities showed the nucleotides had an on rate of about 10^9^*M^−^*^1^*s^−^*^1^ ^13^. Since dNTP has a cytosolic concentration of about 10-15 micro molar^14^, or about 100 fold less than that of ATP (which is at a concentration of 1-1.5 mM^14,15^), the reaction rate k = *k_on_×*[dNTP] = 10^4^ − 10^5^*s^−^*^1^. That is, nucleotides arrive at the arginine ring at a rate of 10^4^ to 10^5^ per second.

Figure 5 shows the potential profile along the pore axis. The potential is computed in the same way as what is shown in Fig. 1, using the APBS software^17^. From the potential profile along the pore axis, it is seen that the barrier height to the interior is about 5.5 KT/e, since nucleotide carries 4 units of negative charges, this amount a barrier height of 22 kT. As a result, the off rate, comparing to the on rate, is reduced by *e*^22^ = 10^9^ fold, to 10^−4^ per second. The rate of binding and reacting with DNA reverse transcriptases may occur even slower^15^. This rate is apparently too slow for DNA synthesis in HIV-1 capsid, which is known to take place at a rate of about 70 nucleotides per minutes, or 1.1 nucleotides per second^19^.

**Figure 5.**
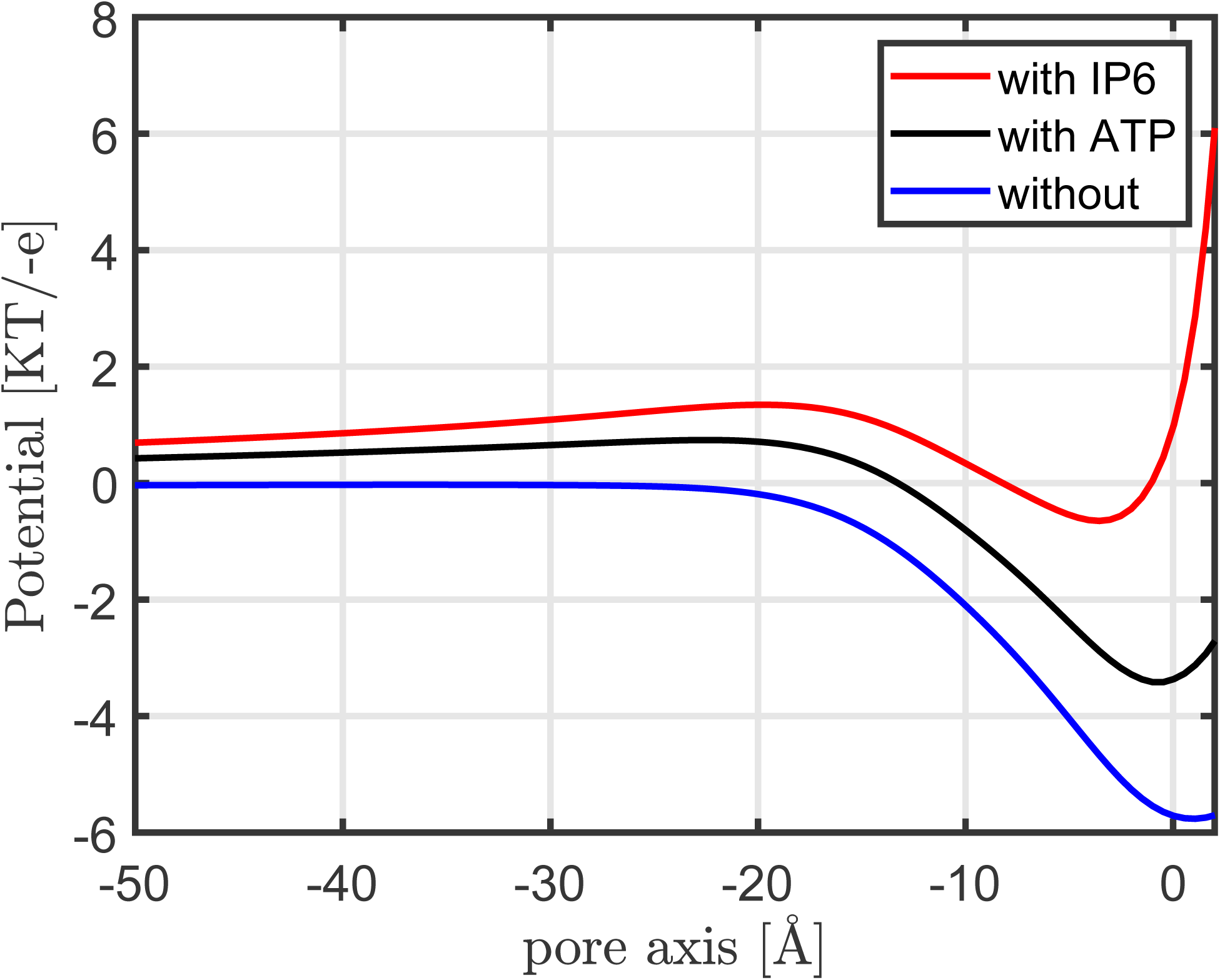
The potential profiles along the pore axis with IP6 (in red) or ATP (in black) binding at the arginine ring, or without. IP6 or ATP binding greatly lowers the potential barrier for dNTP to escape, with IP6 having a greater effect than ATP.

Therefore, a nucleotide must have received outside help for a faster release from the arginine ring into the capsid interior. We hypothesize that the binding of the next nucleotide (dNTP or ATP) or an IP6 should greatly accelerate the release rate.

### 3.5 IP6 facilitates the disassociation of dNTP from the arginine ring

To test the above hypothesis, we recompute the potential within the pore with a nucleotide or an IP6 binding at above the arginine ring using the APBS software (see Fig. 5). As an approximation, IP6 is represented by a sphere of radius 5 Å and with 6 units of negative charge and is place at 5 Å above the arginine ring. (Similarly, the nucleotide is modeled as a sphere of radius 3.5 Å and with 4 units of negative charge and is placed at 3.5 Å above the arginine ring. In this model, there is no distinction between ATP and dNTP).

With the help of IP6, the barrier height to cross before entering the interior of the capsid is reduced from 22 kT to 5 kT with IP6 binding and 16 kT with ATP/dNTP binding (see Figure 5. The rate is thus increased by about *e*^17^ = 10^7^ fold, to 1,000 per second with IP6 binding, which is sufficiently fast for DNA synthesis. With ATP/dNTP binding, the rate is increased by about 10^3^ fold, to 0.1 per second. Since there are hundreds of hexamers on the capsid, the total rate will be further increased by one or two orders, probably making it fast enough for DNA synthesis (which uses 1.1 nucleotides per second^19^).

In summary, the binding of the next nucleotide accelerates the release rate of bound nucleotide from the arginine ring by about a thousand fold. The binding of IP6 has an even greater impact, increasing the release rate by millions of fold.

Figure 6 shows the diffusion process of a nucleotide under the new potential with IP6. The simulation is stopped when the nucleotide reaches 30 Å below the arginine ring. The trajectory of the nucleotide is shown in black. The presence of an IP6 at the arginine ring greatly lows the energy barrier that the nucleotide has to cross in order to enter into the interior of the capsid, as shown in Figure 6.

**Figure 6.**
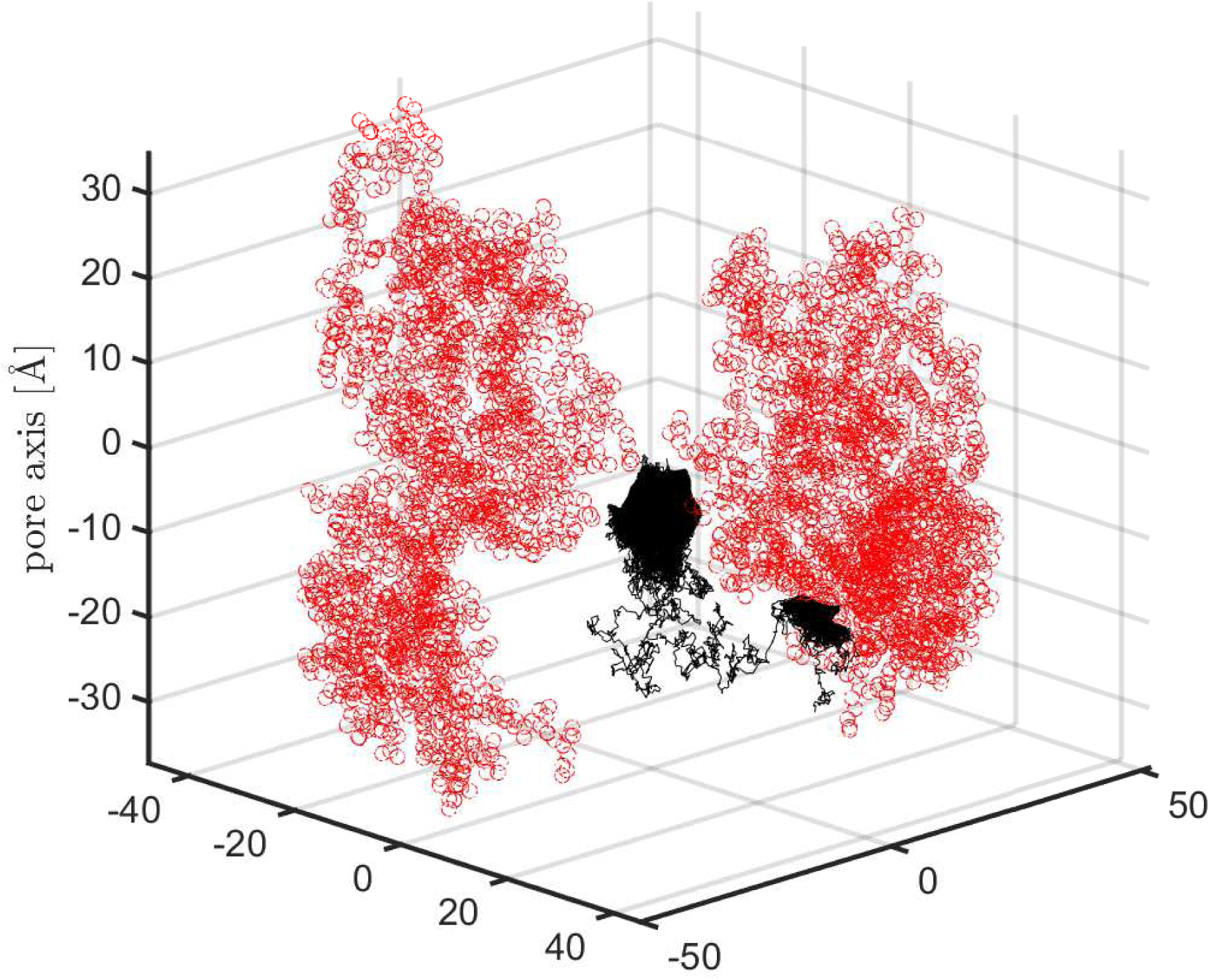
The release of nucleotide in the presence of IP6. The binding of IP6 greatly lowers the barrier height (see Figure 5) that a nucleotide has to cross in order to diffuse into the interior of the capsid. The blue line shows the Brownian trajectory of the nucleotide. The simulation lasted 103,827 steps, or 1.3 *µ*s.

#### Estimate the release time and simulation steps needed

In the following we estimate the amount of time needed for a nucleotide bound at the arginine ring to break away and diffuse to a distance about 30 Å below the arginine ring.

According to the Fick’s second law of diffusion, the approximate time required to reach a distance of *r* is:

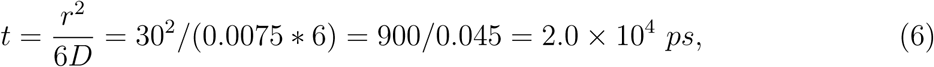

where D = 0.0075 Å^2^*/ps* is the diffusion coefficient of the nucleotide^15^. Because of the steric restriction provided by the hexamers, the downward diffusion should be accelerated by a few folds. On the other hand, the energy barrier height between the bound state at the arginine ring and the capsid interior is about 1.3 kT/(-e) (see Figure 5). Considering the four units of negative charge of the nucleotide, this amounts to a barrier height 1.3 *×* 4 = 5.2 kT. This will slow the diffusion process by *e*^5.2^ = 180 fold. So the total diffusion time will be about 10^4^ *×* 10^2^ = 10^6^ ps, or 1 *µs*. The simulation step size is 12.5ps. Therefore the number of simulation steps needed will be on the order of 10^5^ steps.

The actual simulation steps of the 3-D Brownian simulation shown in Figure 6 is 103,827 steps, which is in good agreement with the above estimation.

#### The dual role of IP6

Therefore, we conclude that IP6 should have a dual role at the arginine ring: it not only serves to stabilize the capsid^14^ but also plays a key role in facilitating dNTP release from their binding at the arginine ring. Even if we take into account of the fact that IP6 competes for arginine ring binding and reduces the on rate of nucleotide binding, its binding at the arginine ring over-compensates the reduction in on rate by increasing the off rate of the nucleotides by millions of fold. As a result, IP6 still significantly accelerates dNTP import. This solves and explains the puzzle why the rate of nucleotide import, instead of being reduced, increased significantly in the presence of IP6 ^14^.

### 3.6 What displaces IP6?

So far we have seen that the binding of the next nucleotide facilitates the disassociation of a dNTP from the arginine ring (Fig. 5). The effectiveness of an IP6 in facilitating the disassociation is even greater, probably due to its more planar geometry that should align better with that of the arginine ring and the extra negative charges it carries. Indeed, IP6 was shown to bind to the ring substantially stronger and stabilize the capsid at much lower concentrations than ATP^14^, and most probably, than dNTP as well. So a natural follow-up question is, once an IP6 is tightly bound to the ring, what helps displace it, even if just temporarily, so that the pore may be used for transporting other cargos such as dNTPs, ATPs, NNRTIs, NRTIs, etc.?

Our hypothesis is that the stochastic Brownian motions of nucleotides (ATP or dNTP) can facilitate the displacement of IP6 from its binding at the arginine ring in the same fashion as they do to the nucleotides, though it will probably take a longer time to displace an IP6 since it has a stronger binding affinity with the arginine ring. To test this hypothesis, we run a Brownian simulation with both a nucleotide (ATP/dNTP) and an IP6. The Brownian motions of the two particles are influenced by the potential field created by the hexamer as shown in Figure 1. The interaction between the nucleotide and the IP6 is through a standard electrostatic interaction using a dielectric constant of 78.54 (Eq. 1). IP6 is modeled as a sphere of radius 5.0 Åfor easy Brownian simulations. It six negative charges however are not placed at the center of the sphere but distributed at the vertices of a regular hexagon horizontally placed within the sphere. The side length of the hexagon is 4.2 Å, the average distance between neighboring phosphate groups within an IP6. Initially, the IP6 is placed at 5 Å above the arginine ring, while the nucleotide is placed at 15 Å below the arginine ring. The simulation terminates when either the nucleotide or IP6 exits from the top or from the bottom (into the interior of the capsid). The simulation lasts nearly 1 million time steps (or 12.5 *µ*s).

Figure 7 shows the trajectories of the nucleotide (in blue) and the IP6 (in black) as a function of time, particularly their coordinates along the pore axis. During the process, both the nucleotide and IP6 seem to make transition between two states. Fig. 8 shows the histogram distributions of the nucleotide and IP6’s displacements along the pore axis. There appear to be two distinct states for both the nucleotide and IP6. For the nucleotide, one state is mostly below the arginine, the other is about 10 Å above the arginine ring. The former clearly corresponds the energy well below the arginine ring in the presence of IP6 as seen in Figure 5. The latter corresponds to an energy well above the ring, again resulting from the net effect of the IP6’s potential and the potential within the pore by the hexamer. Similarly, the potential within the pore by the hexamer and the potential generated by the nucleotide create two energy wells for IP6: one of which is about 5 Å above the ring, which is the binding state for IP6, while the other state is 10 Å above the ring. That both the nucleotide and IP6 have a state at 10 Å above the ring also coincides with the fact that the chamber has the largest cross section at this distance above the ring, as shown in Fig. 1.

**Figure 7.**
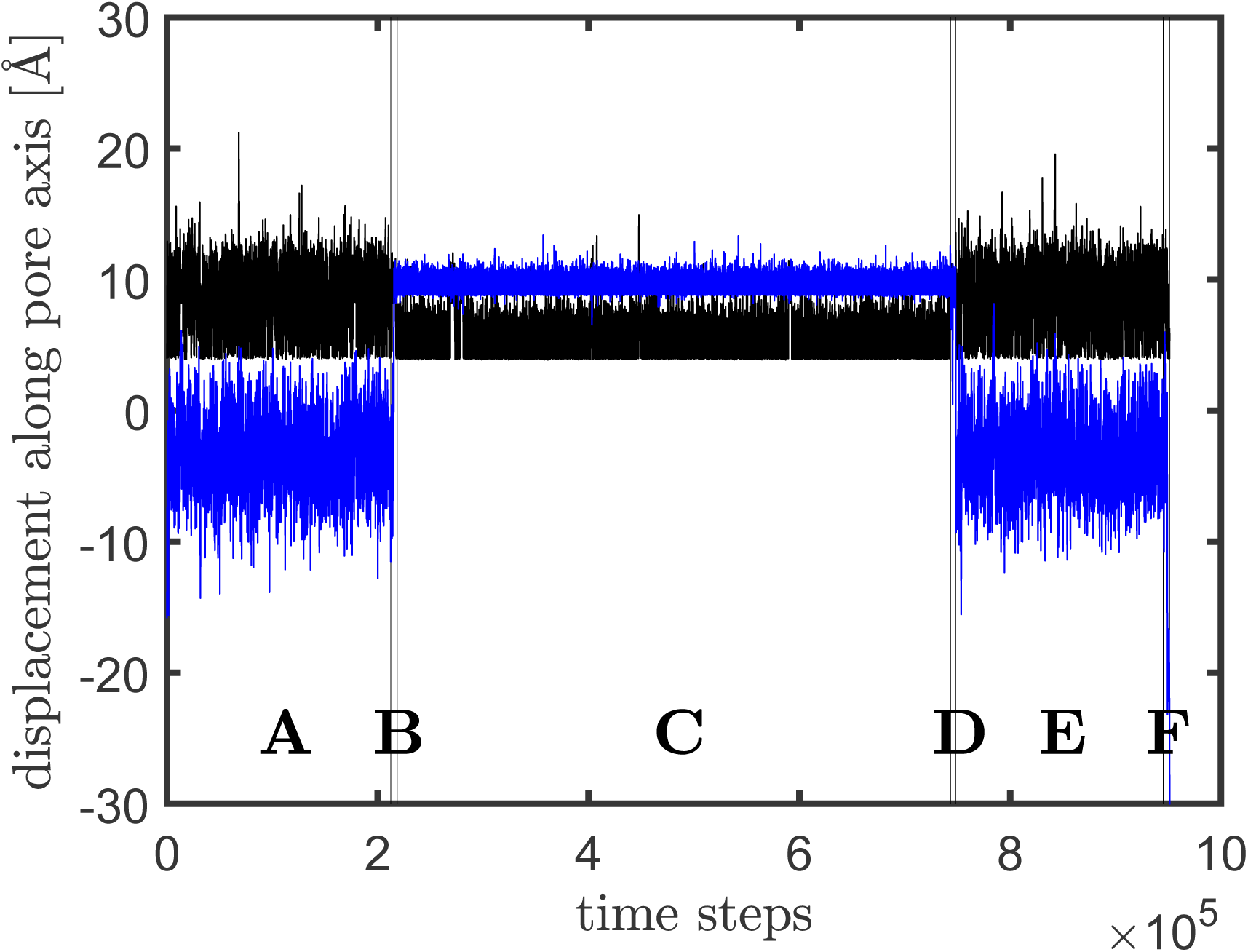
The locations of the nucleotide (blue) and IP6 (black) along the pore axis as a function of time. Each time step is 12.5 ps.

**Figure 8.**
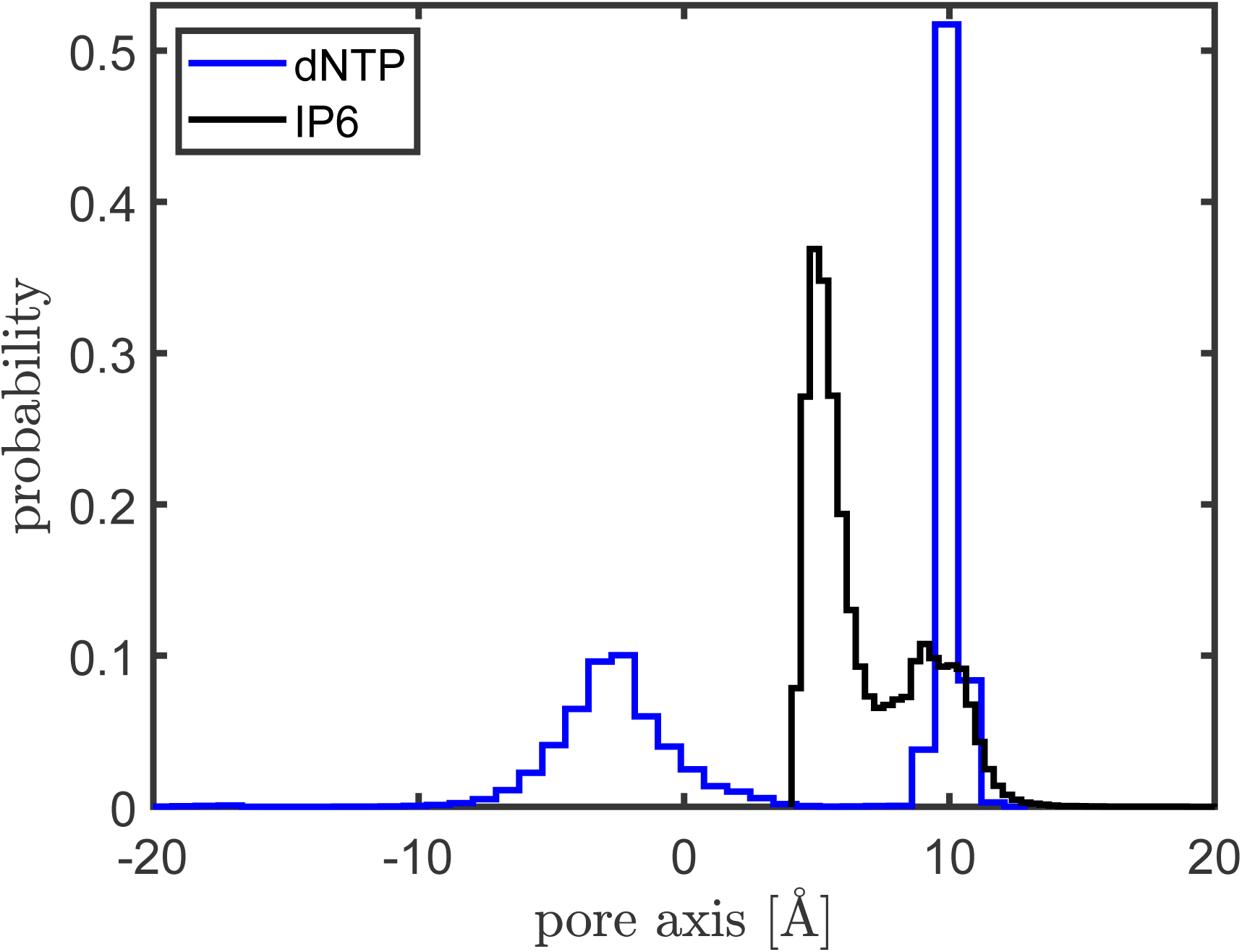
The probability distributions of the nucleotide and IP6 along the pore axis according to one Brownian simulation that lasts about 12 *µ*s. The distributions show that both the nucleotide and IP6 have two states. The nucleotide has a state below the arginine ring (on the negative side of the abscissa axis) and a state above the arginine ring (on the positive side of the abscissa axis). Both states of IP6 are within the chamber above the arginine ring: one is at the binding position near the ring while the other is at about 10 Å above the ring, where the pore has the largest cross section (see Fig. 1).

The time evolution of the trajectories of the nucleotide and IP6 (Figure 7) shows that the two particles go through several distinct stages, as labeled by letters **A**-**F** in the bottom of Figure 7. From the figure we see also that when the nucleotide is at its first state below the ring (stage **A**), IP6 switches between its two states frequently. When the nucleotide makes a transition (stage **B**) from below the ring to above the ring and reaches to its other state above the ring (stage **C**), IP6 occupies nearly exclusively its lower, biding state at the ring. At the end of stage **C**, the nucleotide makes another transition (stage **D**) and moves from above the ring to below the ring, where it stays for an extended period of time (stage **E**). Then it leaves the energy well below the ring again. The difference is this time it moves downwards and enters into the interior of the capsid (stage **F**). Once the nucleotide has moved downward into the interior, IP6 stays exclusively in the binding state right above the ring (see Figure 7, stage **F**, and also Fig. 3).

Fig. 9 shows the corresponding 3-D trajectories of the nucleotide and IP6 at these six stages (stages **A**-**F**). For clarity, only the heavy atoms of two protein chains (chains A and D) of the hexamer are shown and they are represented by red circles. The trajectory of the the nucleotide is in blue while that of IP6 is in black. It is seen that at stage (**A**) the nucleotide indeed stays below the ring. At stage (**B**) it moves up to above the ring. It can do this because at this time IP6 has left its biding state temporarily due to the repulsion by the nucleotide and moved up in the chamber (see Fig 9(B)). Fig 9(C) shows that there is a stage when both particles are in the chamber. As aforementioned, the chamber above the arginine ring (Figure 1) is large and is over 3,000 Å^3^ in volume^13^ and probably can host several nucleotides and/or IP6s at a time. Though our simulations shows only the interplay of two particles (a nucleotide and an IP6), the displacement of the IP6 from the arginine ring by the nucleotide and how they repel each other in the chamber as shown in stage (**C**) probably creates opportunities for other cargos such as other nucleotides, NNRTIs, NRTIs, etc. to utilize the pore for transport. Therefore, though IP6 binds to the arginine more strongly than ATP or dNTP, it does not bind there all the time nor block the transport route.

**Figure 9.**
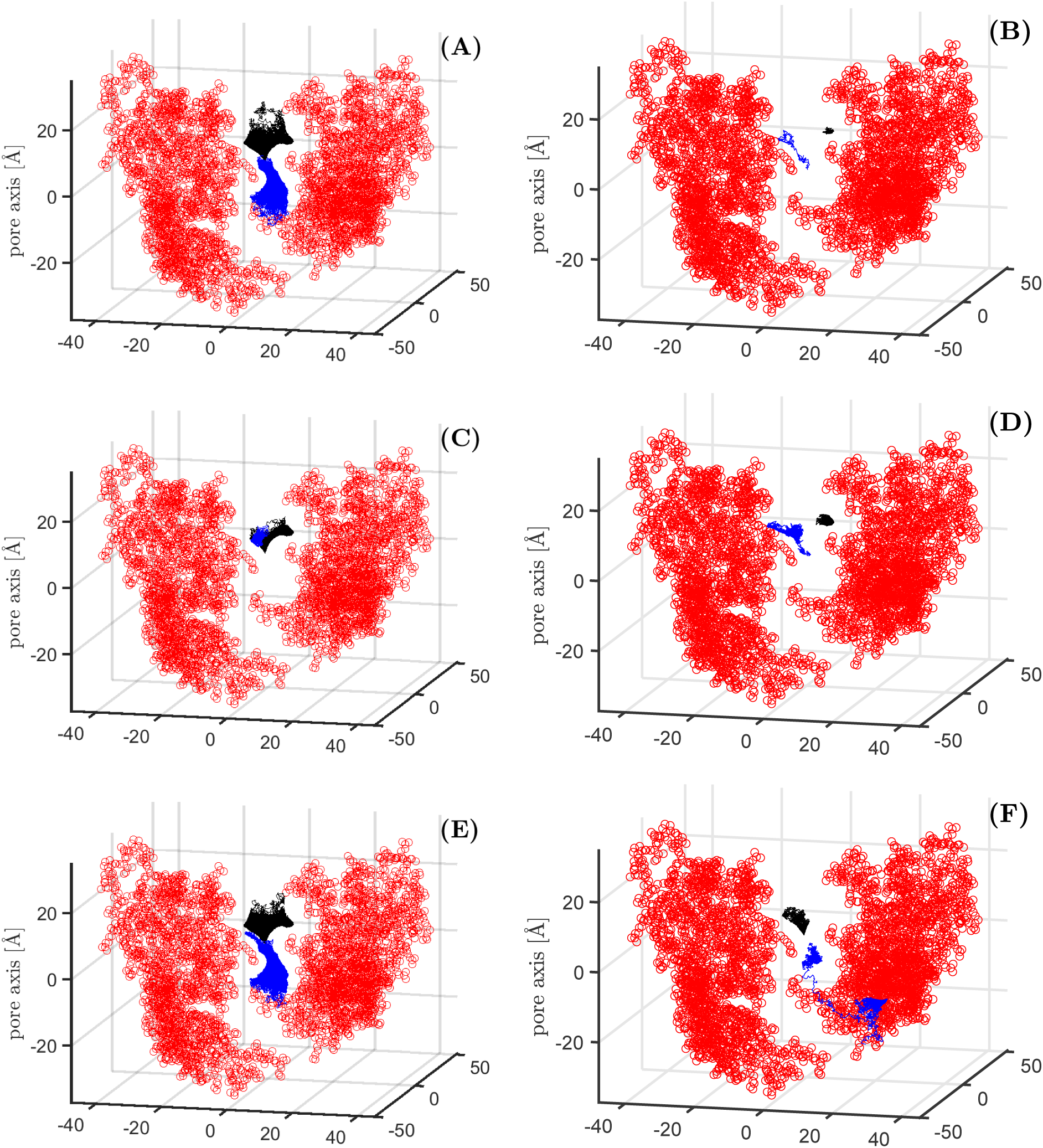
The diffusion of a nucleotide displaces IP6 from its binding at the arginine ring. Panels **A**-**F** shows the Brownian trajectories of the nucleotide (in blue) and IP6 (in black) at different stages (**A**-**F**) of the simulation process as labeled in Fig. 7.

Fig. 9(D) shows a nearly reverse process of stage (B), where the ATP (in blue) makes a transition from above the ring to below the ring. Again IP6 moves up in the chamber at this time to open the way, so that it does not block the narrow hole in center of the arginine ring. Stage (**E**) (Fig. 9) is a extremely similar to (**A**), which is expected. This difference is that at the end of stage (**E**), the nucleotide moves downward and takes a trek into the interior of the capsid.

In the above Brownian simulation, IP6 is modeled as a sphere with a radius of 5.0 Å and a diffusion coefficient of 0.0075 Å^2^/ps. Brownian simulations of non-spherical objects are much more difficult and thus are not considered in the present study. The presence of ions also influence the diffusion behaviors^20^. However, the effects of ions are not considered except when computing the potential field generated by the hexamer using the APBS software. Regarding binding energy, only electrostatics is considered. This is a rough estimate. The actual binding energy may be stronger and the diffusion coefficient may be reduced also after the binding. As a result, the actual process of displacing IP6 may take a longer time. Nevertheless, the principle seen here, that the diffusion of the nucleotides and their interactions with IP6 greatly facilitates the displacement of IP6 or vice versa, should still hold.

## 4 Discussion

In this work, we present Brownian simulations of the diffusion process of deoxynucleoside triphosphates (dNTPs) through the R18 pores of HIV-1 capsid. The simulations provide a clear explanation of how the positive charges at the arginine ring help recruit nucleotides and cause them to arrive at the arginine ring at a near diffusion-limited rate. Our computational analysis reveals also that without outside help, a bound nucleotide would break off from the arginine ring and diffuse into the interior of the capsid at a rate of only 10^−4^ per second, which is far too slow to fuel the reverse DNA synthesis inside the capsid. Therefore, a bound nucleotide must have received outside help for its release from the arginine ring.

### The dual role of IP6 at the arginine ring

Our simulations of the nucleotide diffusion in the presence of IP6 uncovers a novel role of IP6. IP6 plays a key role in the stabilization of the capsid^14^. Specifically, the presence of IP6 increased the viral stability from minutes to hours and ERT efficiency by about a hundred folds^14^. The authors attributed this increase in ERT activity to the increased stability^14^. In this work, we demonstrate that, besides increasing viral stability, IP6 also significantly accelerates the release rate of nucleotides from the arginine ring. The identification of this second role explains a number of puzzling facts that could not be well explained by using the stability theory alone, such as why IP6 or ATP, as a much stronger competitor for arginine ring binding, does not block out ERT activity altogether.

### The sequential binding-induced release mechanism of nucleotide import by HIV-1 capsid

In the absence of IP6, it would take a second incoming nucleotide’s binding to the pore to displace the first bound dNTP into the capsid interior, employing a sequential binding-induced release mechanism, or a nucleotide-induced nucleotide release (NINR) mechanism (see the schematic representation of the process in Scheme 1). The core nature of this mechanism is obviously physical and electrostatic, not through biochemical properties of ATP such as its hydrolysis, but through their charges. Therefore it is expected that other polyanions should have similar effects in increasing ERT efficiency if they are present in the system at a similar concentration. Evidence supporting this hypothesis is that ATP*γ*S, an ATP analog that carries the same charge as ATP, was found to produce a similar increase in ERT efficiency^14^.

The sequential binding-induced release mechanism of HIV-1 capsid provides an effective way to import the nucleotides. It resembles to some extent the calcium-induced calcium release (CICR)^21,22^ mechanism employed in cells to accelerate calcium propagation.

### Why are pores not clogged by IP6?

IP6 binds to the arginine ring substantially stronger than nucleotides and yet it does not block the pore from transporting other molecular cargos. Our simulations explain why this can be the case. Scheme 2 (also Fig. 9) shows how IP6 may be displaced from the arginine ring due to the diffusion and repulsion of nucleotides. Its displacement, even if just temporarily, opens the pore to be used by other cargos. This displacement should be mostly caused by ATP instead of dNTP since the former is about 100 times more abundant than the latter^14^. In other words, the constant flow (diffusion) of ATPs in and out of the capsid should cause IP6 to be displaced from time to time, opening up the pores for other particles to diffuse through.

### Why is ATP’s effect in improving ERT efficiency dose-dependent?

IP6 and ATP were thought to compete with dNTP for binding at the capsid hexamers and it was puzzling why IP6 or ATP binding did not hinder nucleotide import but rather accelerated it. Our work provides an explanation. It shows that IP6, though indeed competes for binding at the R18 pores and reduces the availability of pores for dNTP import and thus its on rate, its binding helps accelerate the off rate of nucleotides by such a large number of folds that it sufficiently offsets the reduction it causes in the availability of pores. ATP, on the other hand, is comparable to dNTP, both having a much weaker binding affinity to the arginine ring than IP6. Therefore, it is expected that ATP’s effect in accelerating dNTP off rate and consequently ERT’s efficiency is dose-dependent (Figure 2c in Ref. 14), since higher ATP concentrations increase the on rate of ATP and thus the release rate (off rate) of dNTP (Scheme 1). It is expected also that when dNTP concentration reaches to a high enough level, it can assist its own release sufficiently (Scheme 1) and the presence of ATP should not make a difference. This indeed was found to be the case: that ATP helps only when the concentrations of dNTPs is below 100 *µ*M (Figure 2f in Ref. 14).

### Why do the capsid pores show no preference for nucleotide-based reverse transcription inhibitors (NRTIs) over non-nucleoside reverse-transcriptase inhibitors (NNRTIs)?

The present work answers yet another puzzle. Mallery et al.^14^ noted that “if pores represent the only entry route into the capsid then NNRTIs might be less potent in an encapsidated assay because they lack a triphosphate and may fail to interact with the R18 ring.” However, they found that the pore were able to import both inhibitors equally well, showing no preference for one over the other^14^. This was puzzling. Our diffusion simulations explain how NNRTIs and NRTIs can share the same pore and have similar rates in passing through the arginine ring and entering into the capsid. The NNRTIs, though without a triphosphate that binds to the R18 ring and as a result having a much lower on rate than NRTIs do, can diffuse thorough the arginine ring nearly effortlessly since they face no high energy barrier in entering into the capsid interior that NRTIs have to cross. As illustrated in Fig. 2, the hexamer pores should be open from time to time for these inhibitors and other small molecules to pass through.

### Uncoating: what triggers the release of IP6?

A key step in the viral life of the HIV-1 is the disassembly of the capsid core, a process also called uncoating. Several theories exist regarding what triggers the process. By applying atomic force microscopy, Rankovic et al. observed that reverse transcription elevated the pressure inside the capsid and proposed it to be the cause for initiating the uncoating process^23^. Mallery et al.^14^, after identifying the key role of IP6 as a pocket factor in stabilizing the capsid, proposed that the release of the IP6 should trigger the uncoating. However, what triggers the IP6 release was not answered. Our simulation results presented here suggest that the IP6 release signal probably originates from within the capsid, similar to that shown in Scheme 2. Once DNA synthesis is completed, the concentration of the dNTP inside the capsid should start to build up. The accumulated concentration might play a role in both increasing the pressure inside the capsid as observed by Rankovic et al.^23^ and helping trigger the release of IP6^14^.

### IP4

There are still some questions left not fully answered. For example, Mallery et al. ^14^ found that while IP6 and IP5 and ATPs accelerate the rate of ERT, IP4 does not. The mechanism proposed in this work (see Schemes 1 and 2) implies that IP4, as a polyanion like the nucleotides or IP6, should accelerate the reverse transcription as well. So why was IP4 not found to accelerate ERT in Mallery et al’s study^14^?

One possible explanation is that the concentration of IP4 was not high enough. IP6 binds to the arginine ring much more strongly than ATP and can stabilize the capsid at a much lower concentration^14^. IP4 is significantly different from IP6, since it carries only four units of negative charges instead of six. Consequently, IP4 may bind and stabilize the capsid in a similar strength to that of ATP. Since ATP’s improvement in ERT efficiency is dose-dependent and is minimal when in a similar concentration to dNTP, we predict that IP4’s improvement in ERT also should be dose-dependent. In order for IP4 to accelerate ERT as noticeably as ATP did, we predict that it would need a similar concentration to that of ATP (about 1 mM), which is much higher than that of IP6 (about 50 *µ*M^24^). And we call for interested experimentalists to carry out this validation.

### The role of hairpins

HIV-1 capsid has hairpins that were thought to undergo iris motions that gate the opening^13^. However, structural analysis of a full capsid structure indicated that the hairpin motions are uncorrelated and stochastic^16^. Seeing the similarity between the hexamer pores on the capsid and the nuclear pore complex (NPC) on a cell’s nucleus, it is tempting to hypothesize that the hairpins should play a similar role to some of what nucleoporins do at the entrance of NPC, namely, making rapid uncorrelated Brownian motions. For NPC, such swinging motions deflect large molecules away from the pore by frequent collisions while small molecules can slip by^25^. The role of the hairpins may be quite similar. Additionally, they may help reduce the escape rate of dNTPs back to cytoplasm^16^.

---

**Scheme 1.**
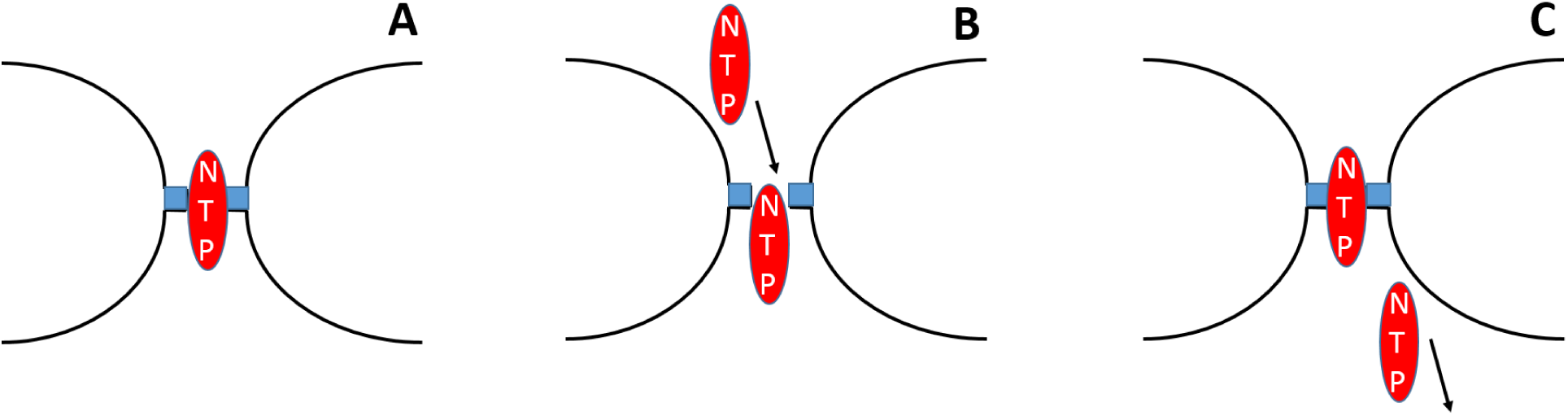
The nucleotide-induced nucleotide release (NINR) mechanism of nucleotide import by HIV-1 capsid. The incoming of next nucleotide (labeled as NTP) facilitates the displacement of the first nucleotide (also labeled as NTP) from its tight binding at the arginine ring, as shown in panels **A**-**C**. The process efficiently produces a constant flux of nucleotides into the capsid.

**Scheme 2.**
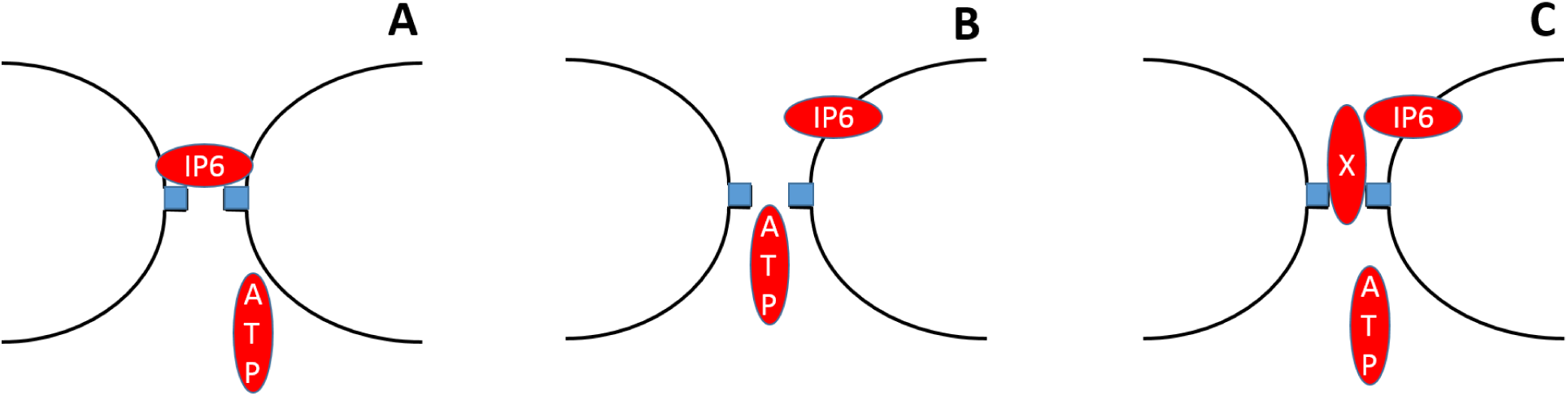
The displacement of IP6 and how it opens up the pores for other cargos. The diffusion of nucleotides (most likely ATPs) (**A**) and their repulsion towards IP6 displace IP6 from its binding at the arginine ring (**B**) and open a way for other cargos (such as dNTP, NNRTI, NRTI, etc. labeled as X in the figure) to use the pore for transport (**C**).

## References

[1] F. Barre-Sinoussi, J. C. Chermann, F. Rey, M. T. Nugeyre, S. Chamaret, J. Gruest, C. Dauguet, C. Axler-Blin, F. Vezinet-Brun, C. Rouzioux, W. Rozenbaum, and L. Montagnier. Isolation of a t-lymphotropic retrovirus from a patient at risk for acquired immune deficiency syndrome (aids). Science, 220(4599):868–71, 1983.

[2] F. Barre-Sinoussi, A. L. Ross, and J. F. Delfraissy. Past, present and future: 30 years of hiv research. Nat Rev Microbiol, 11(12):877–83, 2013.

[3] M. A. Fischl, D. D. Richman, M. H. Grieco, M. S. Gottlieb, P. A. Volberding, O. L. Laskin, J. M. Leedom, J. E. Groopman, D. Mildvan, R. T. Schooley, and et al. The efficacy of azidothymidine (azt) in the treatment of patients with aids and aids-related complex. a double-blind, placebo-controlled trial. N Engl J Med, 317(4):185–91, 1987.

[4] E. J. Arts and D. J. Hazuda. Hiv-1 antiretroviral drug therapy. Cold Spring Harbor Perspectives in Medicine, 2(4), 2012.

[5] G. Zhao, J. R. Perilla, E. L. Yufenyuy, X. Meng, B. Chen, J. Ning, J. Ahn, A. M. Gronenborn, K. Schulten, C. Aiken, and P. Zhang. Mature HIV-1 capsid structure by cryo-electron microscopy and all-atom molecular dynamics. Nature, 497:643–646, 2013.

[6] E. M. Campbell and T. J. Hope. Hiv-1 capsid: the multifaceted key player in hiv-1 infection. Nat Rev Microbiol, 13(8):471–83, 2015.

[7] D. Gao, J. Wu, Y. T. Wu, F. Du, C. Aroh, N. Yan, L. Sun, and Z. J. Chen. Cyclic gmp-amp synthase is an innate immune sensor of hiv and other retroviruses. Science, 341(6148):903–6, 2013.

[8] N. Yan, A. D. Regalado-Magdos, B. Stiggelbout, M. A. Lee-Kirsch, and J. Lieberman. The cytosolic exonuclease trex1 inhibits the innate immune response to human immunodeficiency virus type 1. Nat Immunol, 11(11):1005–13, 2010.

[9] A. J. Price, D. A. Jacques, W. A. McEwan, A. J. Fletcher, S. Essig, J. W. Chin, U. D. Halambage, C. Aiken, and L. C. James. Host cofactors and pharmacologic ligands share an essential interface in hiv-1 capsid that is lost upon disassembly. PLoS Pathog, 10(10):e1004459, 2014.

[10] S. K. Carnes, J. H. Sheehan, and C. Aiken. Inhibitors of the hiv-1 capsid, a target of opportunity. Curr Opin HIV AIDS, 13(4):359–365, 2018.

[11] S. Thenin-Houssier and S. T. Valente. Hiv-1 capsid inhibitors as antiretroviral agents. Current Hiv Research, 14(3):270–282, 2016.

[12] J. Zheng, S. R. Yant, Ahmadyar S., T. Y. Chan, A. Chiu, Tomas Cihlar, John O. Link, Bing Lu, Judy W. Mwangi, William Rowe, Scott D. Schroeder, George J. Stepan, Kelly Wei Wang, Raju Subramanian, and Winston C. Tse. Gs-6207: A novel, potent and selective first-in-class inhibitor of hiv 1 capsid function displays nonclinical pharmacokinetics supporting long acting potential in humans. In IDWeek.

[13] D. A. Jacques, W. A. McEwan, L. Hilditch, A. J. Price, G. J. Towers, and L. C. James. HIV-1 uses dynamic capsid pores to import nucleotides and fuel encapsidated DNA synthesis. Nature, 536:349–353, 2016.

[14] D. L. Mallery, C. L. Marquez, W. A. McEwan, C. F. Dickson, D. A. Jacques, M. Anandapadamanaban, K. Bichel, G. J. Towers, A. Saiardi, T. Bocking, and L. C. James. Ip6 is an hiv pocket factor that prevents capsid collapse and promotes dna synthesis. Elife, 7, 2018.

[15] N. N. Batada, K. D. Westover, D. A. Bushnell, M. Levitt, and R. D. Kornberg. Diffusion of nucleoside triphosphates and role of the entry site to the rna polymerase ii active center. Proc Natl Acad Sci U S A, 101(50):17361–4, 2004.

[16] G. Song. Structure-based insights into the mechanism of nucleotide import by hiv-1 capsid. Journal of Structural Biology, 2019.

[17] N. A. Baker, D. Sept, S. Joseph, M. J. Holst, and J. A. McCammon. Electrostatics of nanosystems: application to microtubules and the ribosome. Proc Natl Acad Sci U S A, 98(18):10037–41, 2001.

[18] T. J. Dolinsky, J. E. Nielsen, J. A. McCammon, and N. A. Baker. Pdb2pqr: an automated pipeline for the setup of poisson-boltzmann electrostatics calculations. Nucleic Acids Res, 32(Web Server issue):W665–7, 2004.

[19] D. C. Thomas, Y. A. Voronin, G. N. Nikolenko, J. Chen, W. S. Hu, and V. K. Pathak. Determination of the ex vivo rates of human immunodeficiency virus type 1 reverse transcription by using novel strand-specific amplification analysis. J Virol, 81(9):4798–807, 2007.

[20] W. Stillwell and H. C. Winter. The stimulation of diffusion of adenine nucleotides across bimolecular lipid membranes by divalent metal ions. Biochem Biophys Res Commun, 56(3):617–22, 1974.

[21] H. L. Roderick, M. J. Berridge, and M. D. Bootman. Calcium-induced calcium release. Curr Biol, 13(11):R425, 2003.

[22] A. Hernandez-Cruz, A. L. Escobar, and N. Jimenez. Ca(2+)-induced ca2+ release phenomena in mammalian sympathetic neurons are critically dependent on the rate of rise of trigger ca2+. J Gen Physiol, 109(2):147–67, 1997.

[23] S. Rankovic, J. Varadarajan, R. Ramalho, C. Aiken, and I. Rousso. Reverse transcription mechanically initiates hiv-1 capsid disassembly. J Virol, 91(12), 2017.

[24] N. Veiga, J. Torres, S. Dominguez, A. Mederos, R. F. Irvine, A. Diaz, and C. Kremer. The behaviour of myo-inositol hexakisphosphate in the presence of magnesium(ii) and calcium(ii): protein-free soluble insp6 is limited to 49 microm under cytosolic/nuclear conditions. J Inorg Biochem, 100(11):1800–10, 2006.

[25] M. P. Rout, J. D. Aitchison, A. Suprapto, K. Hjertaas, Y. Zhao, and B. T. Chait. The yeast nuclear pore complex: composition, architecture, and transport mechanism. J Cell Biol, 148(4):635–51, 2000.

